# Hamstrings muscle dynamics during the Nordic hamstring exercise and high-speed running

**DOI:** 10.1101/2025.09.16.676401

**Authors:** Nicos Haralabidis, Kristen Steudel, Reed Gurchiek, Jennifer Hicks, Scott Delp

## Abstract

**Background:** The Nordic hamstring exercise (NHE) and high-speed running are widely used training modalities to prevent hamstring strain injuries, yet the differences in the muscle lengths, forces, work, and power between these training modalities remain unclear. This study thus compared the dynamics of the most injured hamstrings muscle, biceps femoris long head (BFLH), for 14 participants (8 male and 6 female) performing the NHE and running between 4 and 8 m/s.

**Methods:** We used motion capture experiments and musculoskeletal simulation to quantify muscle fiber lengths and velocities, and muscle force, work, and power during the NHE and running.

**Results:** Our results show that peak muscle forces are greater during high-speed running (7.5 to 8 m/s) than the NHE, and that high-speed running also features longer muscle fiber lengths and higher muscle fiber lengthening velocities (*p* < 0.05). Negative muscle work was significantly greater during the NHE compared to running at all speeds (*p* < 0.001) because of the greater change in muscle fiber lengths during the NHE (*p* < 0.001). In contrast, peak negative muscle power was significantly lower during the NHE compared to running at 5 m/s and above (*p* < 0.01).

**Conclusion:** Our analysis reveals dramatic differences in the biomechanical demands of the NHE and running on the hamstrings muscles. Our results suggest that the two training modalities together provide complementary biomechanical stimuli to promote favorable BFLH injury prevention adaptations.

## 1. Introduction

Hamstring strain injuries account for 17-24% of all injuries in sports that feature high-speed running (e.g., soccer, rugby union, and track and field),^1–3^ and they predominantly affect the biceps femoris long head (BFLH).^4,5^ These injuries have a 13-18% recurrence rate^1,6^ and may prevent an athlete from regaining their pre-injury performance level after they return to play.^6^ Each injury can result in 13-90 days lost from training and competition,^6,7^ which has negative performance^8^ and economic^9^ consequences.

The hamstrings are most susceptible to injury during the late swing phase of the running gait cycle.^10^ During the late swing phase, the hamstrings are highly active, produce high force, reach peak length, and absorb energy at a high rate to arrest the forward motion of the thigh and shank.^11–14^ Increased eccentric strength and fascicle length of the hamstrings are considered favorable adaptations to prevent hamstring strain injuries.^15^ An understanding of the mechanical loads experienced by the hamstrings during various forms of training could be used to prescribe training interventions suited to achieve the desired adaptations.

The Nordic hamstring exercise (NHE) and high-speed running are widely used training modalities to promote favorable adaptations and reduce injury risk.^16–20^ Both have been found to increase hamstrings strength^16–19,21^ and lengthen the BFLH fascicles.^16–18,20,21^ Furthermore, training programs featuring the NHE have been found to reduce hamstring strain injuries,^22,23^ and training programs featuring high-speed running are associated with reduced risk of lower limb injuries.^24^

Despite the favorable hamstrings adaptations from NHE training together with the reduction in injuries from training programs including the NHE, there has been criticism regarding the specificity, dynamic correspondence, and transference of the NHE to the demands placed on the hamstrings during running.^15,25^ Conversely, high-speed running is specific, transferable, and recommended for hamstring injury prevention, as regular exposure may offer protection.^24,26^ A better understanding of the type and intensity of the mechanical loads present in the hamstrings during these two exercises is needed to optimally prescribe exercise programming.^27^

The purpose of our study was to compare the BFLH muscle dynamics while performing the NHE and running across a range of speeds. Experimental measurements and musculoskeletal simulation enabled us to quantify muscle fiber lengths and velocities, and muscle force, work, and power during the NHE and while running at speeds ranging from 4 to 8 m/s. We compared these muscle loading features to examine the differences between these two common training modalities with unprecedented detail. All simulations and experimental data are publicly shared to enable others to reproduce and build on this work.

## 2. Methods

### 2.1. Participants

Fourteen participants (8 males and 6 females; age: 26 ± 5 years; mass: 71.3 ± 11.1 kg; height: 1.75 ± 0.08 m) took part in this study. All participants had a sporting background (e.g., football, soccer, or track and field) and were injury-free during their participation. Ethical approval for the study protocol was granted by the Stanford University Institutional Review Board (IRB-67713), and all participants gave their written informed consent prior to taking part in the study.

### 2.2. Experimental protocol

Each participant completed a top running speed assessment to determine their eligibility to participate. The assessment took place at an athletics track and consisted of three maximal effort 60 m sprints from a stationary starting position. Participants completed a self-led warm up prior to the sprints, and they rested for at least 5 minutes between sprints. We used timing gates (Dashr Timing System, NE, USA) to measure 5 m splits between 35-40, 40-45, and 45-50 m, and calculated each participant’s top running speed using their best single split time from across the three sprints. Participants had to reach a 95% top running speed of ≥ 7 m/s if female or ≥ 7.5 m/s if male to be eligible to complete the study. We initially recruited 17 participants, and 14 met the running speed criterion (mean top running speed: 8.14 ± 0.50 m/s). Participants that met the running speed threshold completed a self-supervised NHE training program for at least four weeks (two sets of six repetitions twice per week), regardless of whether they had prior NHE training, before completing an in-lab biomechanical data collection session.

For the in-lab session, participants first ran at a range of speeds and then performed NHE repetitions while they underwent biomechanical assessment. Participants completed their own warm-up at the beginning of the session and were allowed to rest between trials until they felt ready to continue. Participants performed running trials once at each speed on an instrumented treadmill (Bertec Corp., Columbus, OH, USA). The speed of the first running trial was 4 m/s.

For each subsequent trial, the speed was increased by 1 m/s up to 7 m/s, after which the speed increased by 0.5 m/s increments. The speed of the final running trial did not exceed 95% of each participant’s top running speed. Participants wore a safety harness that was suspended from ceiling beams. The cord attached to the harness and ceiling beams was sufficiently slack to prevent interfering with the participants’ natural running mechanics.

Participants then performed five NHE repetitions on a pair of custom-made wooden boxes placed over in-ground force plates. For each NHE repetition, participants kneeled with one knee on each box, and their ankles were secured to the boxes with Velcro straps. The boxes were each weighed down using two 20 kg weightlifting plates. We instructed participants to complete each NHE repetition as slowly as possible^17^ and to lower their bodies until they could no longer hold themselves up.

### 2.3. Data collection

We measured ground reaction forces at 2000 Hz using an instrumented treadmill for the running trials and two in-ground force plates (Bertec Corp., Columbus, OH, USA) for the NHE repetitions. For the NHE repetitions, we also measured ankle strap forces at 2000 Hz using two uniaxial load cells (HT Sensor Technology, Shaanxi, China). We captured marker positions at 200 Hz using an optical motion capture system (Motion Analysis, Cortex, CA, USA). We attached 52 retro-reflective markers to the surface of each participant’s skin and shoes (see Supplementary Materials for marker locations).

We recorded surface electromyography (EMG) signals from 13 muscles (dominant limb: vastus lateralis, vastus medialis, rectus femoris, medial gastrocnemius, lateral gastrocnemius, soleus, tibialis anterior, BFLH, and semitendinosus; non-dominant limb: vastus lateralis, medial gastrocnemius, soleus, and BFLH) at 2000 Hz using a wireless system (Avanti, Delsys Inc., Boston, MA, USA). We adhered an electrode to each muscle belly after shaving and cleaning the skin and further secured each electrode with medical tape and wraps.

### 2.4. Simulation workflow

To calculate muscle dynamics, we performed EMG-informed inverse dynamics simulations using a direct collocation optimal control approach. For each participant, we simulated three flight phases at each running speed and three NHE repetitions. We processed the motion capture data to determine muscle-tendon kinematics, moment arms, and joint moments (Figure 1A), and together with the EMG data, these data were inputs to perform inverse simulations. Our simulation workflow then consisted of two steps. First, we performed a flight phase simulation at each participant’s top running speed that solved for muscle dynamics while simultaneously calibrating the musculoskeletal model’s optimal fiber lengths, tendon slack lengths, and maximal isometric forces (Figure 1B). Second, we used the calibrated musculoskeletal model to solve muscle dynamics for the remaining trials (Figure 1C).

**Figure 1.**
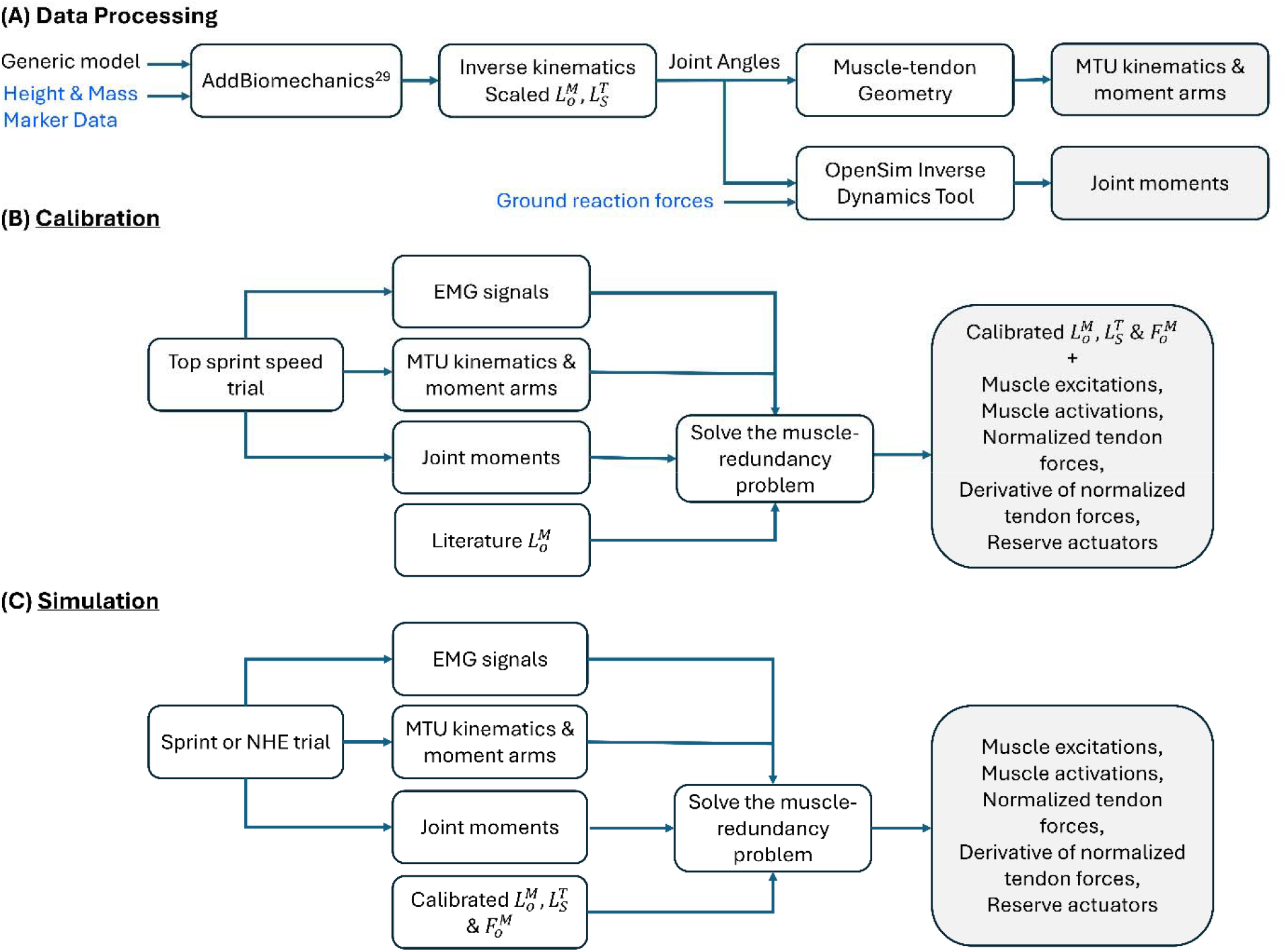
(A) Data processing, (B) model calibration, and (C) simulation workflow (MTU: muscle-tendon unit; EMG: electromyography; 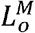:optimal fiber length; 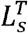: tendon slack length;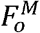:maximal isometric force).

### 2.5. Data processing

We adapted the generic musculoskeletal model of Lai et al.^28^ for this study. For each participant, we used AddBiomechanics (Stanford University, CA, USA)^29^ to simultaneously scale the generic model and calculate joint kinematics based on the marker trajectories from all their running trials and NHE repetitions. We low-pass filtered the joint kinematics for the running trials and NHE repetitions using a fourth-order Butterworth filter at 10 and 4 Hz, respectively.

For the running trials, we analyzed dynamics during the flight phase within which the BFLH reaches peak fiber length and force.^11–13^ For each flight phase, we identified foot-off and foot-strike for the ipsilateral and contralateral limbs, respectively, by applying a 40 N threshold to the filtered vertical ground reaction force (50 Hz cut-off; 4th order low-pass Butterworth filter).

For the NHE repetitions, we analyzed the dynamics until the end of the free fall period when a participant’s hands touched the ground.^30^ An NHE repetition was defined to begin when either of the filtered ankle strap forces (4 Hz cut-off; 4th order low-pass Butterworth filter) first exceeded 50 N. The NHE repetition ended when the vertical velocity of either medial styloid marker first crossed zero.^30^ If neither medial styloid marker had a vertical velocity that crossed zero, the end was defined as the time point when the vertical velocity of either marker was closest to zero, following the peak downward vertical velocity.

We used the OpenSim API (version 4.4; Stanford University, CA, USA)^31^ in MATLAB (2024b; MathWorks Inc., Natick, MA, USA) to calculate net joint moments, muscle-tendon kinematics, and moment arms. For the NHE repetitions, we applied the measured ground reaction forces for each limb to the tibia bodies for inverse dynamics analysis.

Our simulations included an objective term that rewarded muscle excitations that tracked measured EMG signals. We band-pass filtered the EMG data at 30-500 Hz, rectified, and then low-pass filtered at 6 and 12 Hz for the NHE repetitions and running trials, respectively. Filtering was performed using a fourth-order zero-lag Butterworth filter. We normalized each processed EMG signal to the maximum processed EMG value of a given muscle from across all the running trials (including both the flight and stance phases) and NHE repetitions. We then calculated average processed EMG signals for nine muscles per limb. For running, EMG signals were averaged across six flight phases, featuring three flight phases with each of the dominant and non-dominant limbs in late swing. For the NHE, EMG signals were averaged across all three analyzed trials. Four EMG signals were measured on the dominant and non-dominant limbs, and we used their average for both limbs. Five EMG signals were measured only on the dominant limb, and we used this average for both limbs. During processing, we noticed that the signal quality of one to three EMG signals was poor and unusable for four participants, and we used the cohort’s average EMG data to fill in the unusable EMG data for the relevant muscles.

### 2.6. Inverse simulations

Given the processed muscle-tendon kinematics, moment arms, joint moments (Figure 1A), and EMG signals, we then calibrated the musculoskeletal model (Figure 1B) and simulated muscle dynamics (Figure 1C) during the experimental trials using an approach similar to the Muscle Redundancy Solver.^32^ The objective function for both calibration and non-calibration simulations consisted of four weighted terms: sum of squared activations, sum of squared passive force multipliers, sum of squared differences between excitations and EMG signals, sum of squared derivative of tendon forces, and sum of squared reserve joint moment actuators. In total, 13 EMG signals per limb were tracked. We tracked nine processed EMG signals per limb that directly mapped to the corresponding muscle excitations in our model. Additionally, we tracked a subset of the processed EMG signals for each limb that corresponded to adjacent muscle excitations in our model:^33^ average of vastus medialis and vastus lateralis to vastus intermedius, BFLH to biceps femoris short head, and semitendinosus to semimembranosus. For all simulations, we assigned the highest weights in the objective function to minimize the reserve joint moment actuators and EMG tracking error. For the calibration simulations, the objective function also included squared, normalized deviations of optimal fiber length, tendon slack length, and maximal isometric force from their nominal values as additional, weighted terms. We set the bounds of the optimal fiber lengths according to experimentally reported values,^34^ whereas we set the bounds of the tendon slack lengths and maximal isometric forces based on prior literature.^30,35,36^ Further details on the objective functions, constraints, and musculoskeletal model can be found in the Supplementary Materials.

Each simulation was formulated as an optimal control problem. We transcribed each optimal control problem into a nonlinear programming problem using trapezoidal collocation with 50 and 150 mesh intervals for the flight phase and NHE trials, respectively. We formulated all simulations in MATLAB using CasADi,^37^ and solved them using IPOPT^38^ with an adaptive barrier parameter strategy and convergence tolerance of 10^-3^.

To verify the fidelity of the simulated muscle excitations we compared them to measured EMG signals (Figure 2). This revealed that the patterns of the muscle excitations for the BFLH and semitendinosus reproduced the salient features of the measured EMG signals for both running and the NHE with a maximum root mean square error (RMSE) of 0.05 (Figure 2). The patterns of the muscle excitations for the rectus femoris and lateral gastrocnemius also captured the characteristics of the measured EMG signals except for the underestimation of lateral gastrocnemius excitation during running at 8 m/s. The remaining muscle excitations for which we tracked EMG data can be found in the Supplementary Materials (Figure S1).

**Figure 2.**
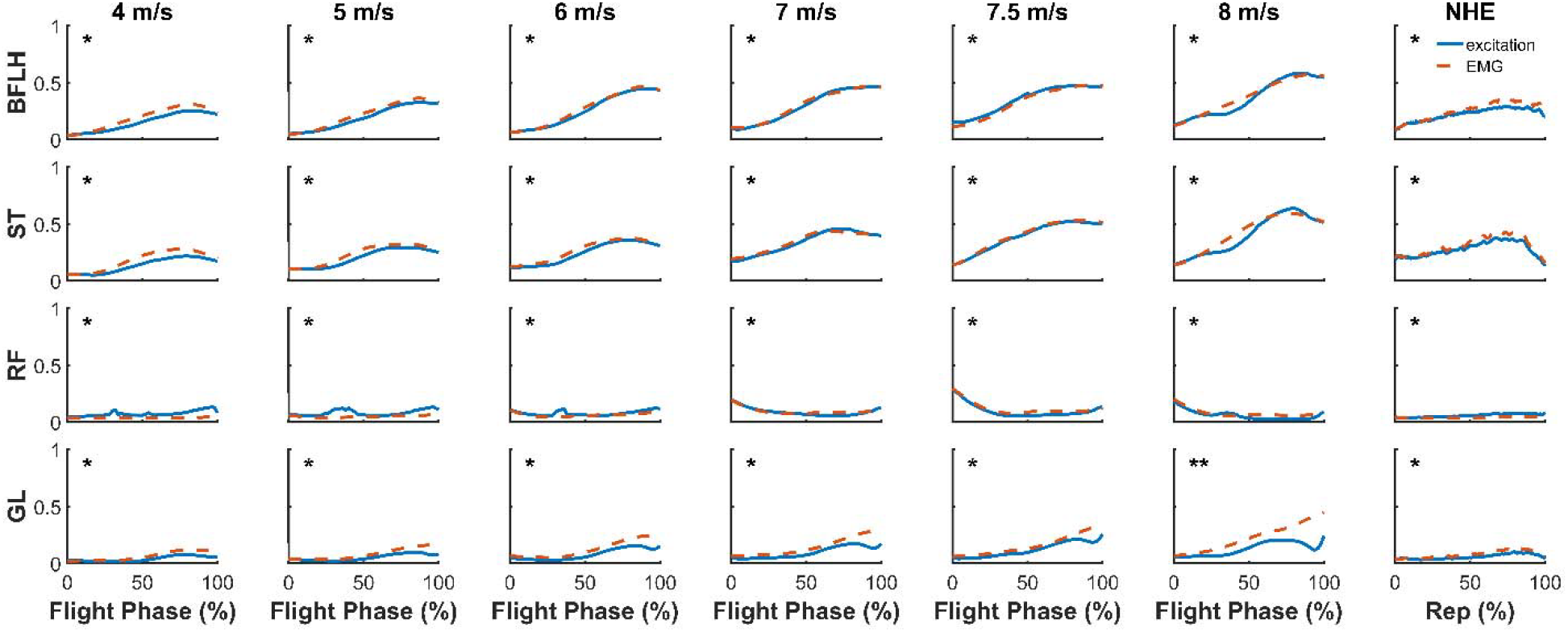
Simulated muscle excitations (solid blue line) and processed EMG signals (dashed red line) during the flight phase of high-speed running at different speeds and during the NHE. Each curve is the mean from across all participants’ trials. Asterisks indicate the magnitude of the root mean square error (RMSE) between the mean muscle excitations and processed EMG signals: * = RMSE of ≤ 0.06 and ** = 0.06 < RMSE ≤ 0.12. The figure includes a subset of key lower-limb muscles, including the biceps femoris long head (BFLH), semitendinosus (ST), rectus femoris (RF), and lateral gastrocnemius (GL).

### 2.7. Outcome variables

We focused on the BFLH dynamics for the forward limb during the flight phase and the same single limb for the NHE. The outcome variables we compared between high-speed running and the NHE included peak muscle force, peak normalized fiber length, normalized fiber length excursion (the peak change in normalized fiber length from the beginning of the flight phase or NHE repetition), peak normalized fiber lengthening velocity, peak negative muscle-tendon unit power, and negative muscle-tendon unit work. We calculated muscle-tendon unit power as the product of tendon force and muscle-tendon unit velocity. We integrated negative muscle-tendon unit power with respect to time to calculate negative muscle-tendon unit work, and negative work represented power absorption.

### 2.8. Statistical analysis

We performed statistical analyses using RStudio (version 4.4.2).^39^ For each outcome variable we fit a linear mixed effects model to the running speeds using the lme4 package.^40^ For the NHE, we calculated the sample mean of all the repetitions for each outcome variable. We used the emmeans package^41^ to compare estimated marginal means of each running speed to the NHE sample mean using a t-test with a Bonferroni correction.

## 3. Results

The peak normalized fiber length of the BFLH during the NHE was significantly shorter compared to running at 6 m/s and above (*p* < 0.05) (Figure 3A), whereas the normalized fiber excursion was significantly greater for the NHE compared to the flight phase of all running speeds (*p* < 0.001). The peak normalized fiber lengthening velocity of the BFLH during the NHE was significantly lower compared to running at all speeds (*p* < 0.001) (Figure 3B).

**Figure 3.**
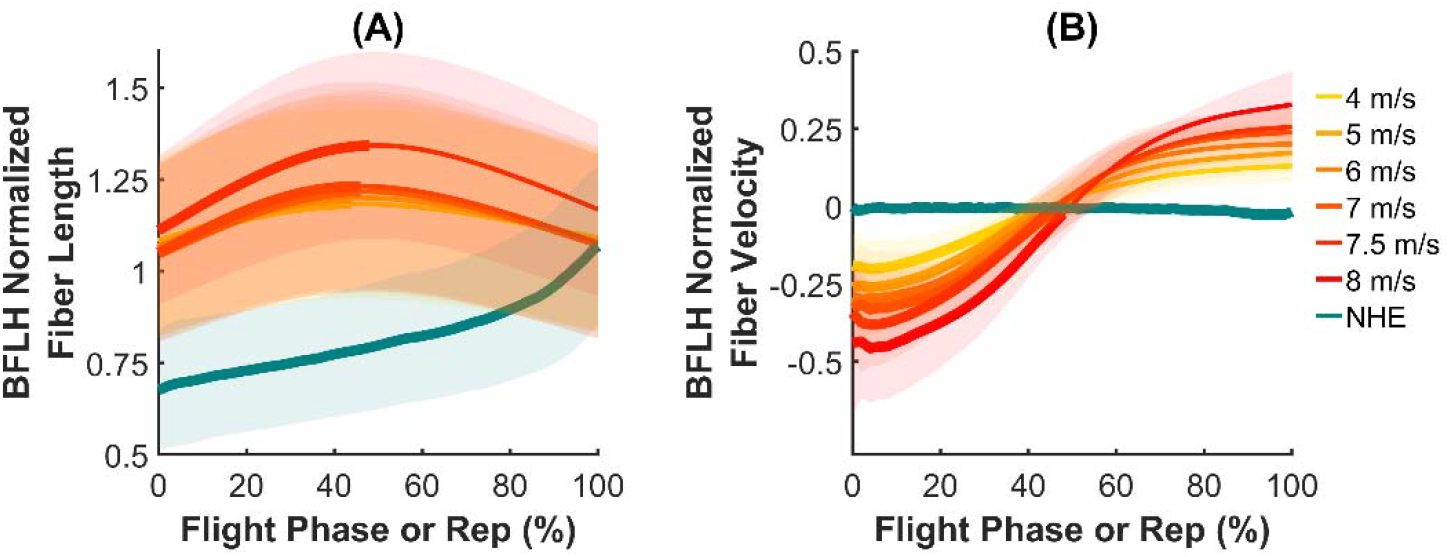
Biceps femoris long head (BFLH) normalized fiber length 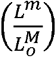 (A) and BFLH normalized fiber velocity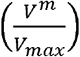(negative velocity indicates fiber lengthening) (B) during the flight phase of high-speed running at different speeds and the NHE. Solid lines and shaded areas indicate the group mean ± one standard deviation from across all participants’ trials. The region over which the BFLH fiber is lengthening while running is indicated by a thick line; the thinner line is the region over which the BFLH is shortening.

The peak BFLH force was significantly greater for the NHE compared to the flight phase of running at 4 m/s (*p* < 0.01) (Figure 4A); however, the peak BFLH force was significantly lower for the NHE compared to the flight phase of running at 7.5 m/s and above (*p* < 0.05). There was no difference in peak BFLH force between the NHE and running at 5, 6, and 7 m/s. The peak negative muscle-tendon unit power during the NHE was significantly lower compared to the flight phase of running at 5 m/s and above (*p* < 0.01) (Figure 4B). Conversely, the negative muscle-tendon unit work was significantly greater during the NHE compared to the flight phase of running at all speeds (*p* < 0.001) (Figure 4C).

**Figure 4.**
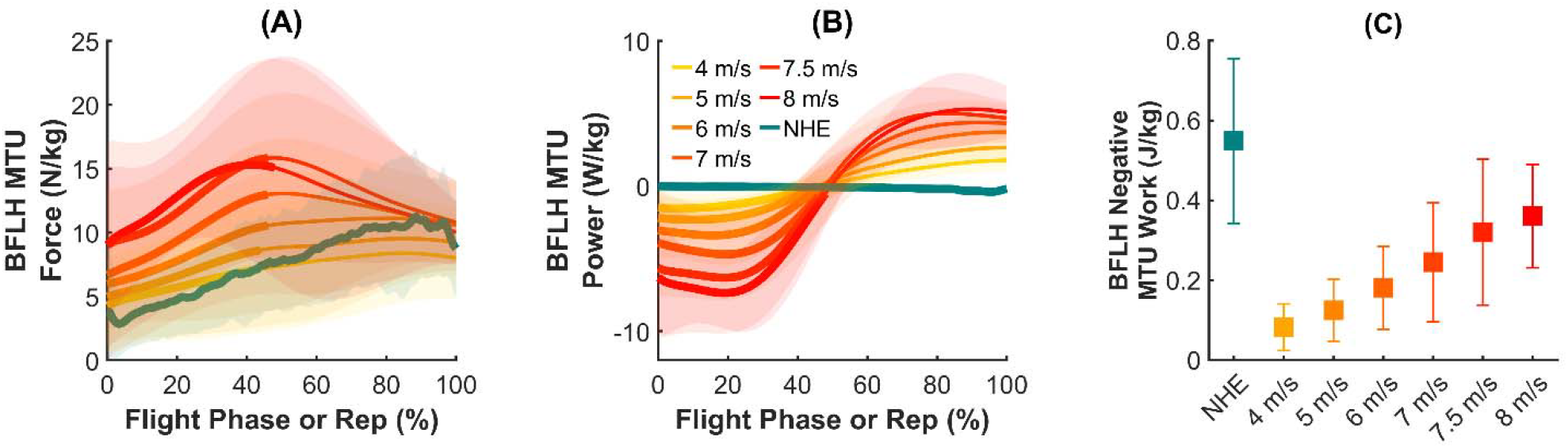
Biceps femoris long head (BFLH) muscle-tendon unit (MTU) force (A), BFLH MTU power (negative power indicates fiber lengthening) (B), and BFLH negative MTU work (C) during the flight phase of high-speed running at different speeds and the NHE. Solid lines and shaded areas indicate the group mean ± one standard deviation from across all participants’ trials. The region over which the BFLH fiber is lengthening while running is indicated by a thick line (A & B); the thinner line is the region over which the BFLH is shortening.

### 4. Discussion

The purpose of this study was to compare BFLH muscle dynamics during the NHE with those during running across a range of speeds. While peak muscle forces were similar between the NHE and running at 5, 6, and 7 m/s (i.e., no significant differences), the peak forces were significantly lower during the NHE compared to running at 7.5 m/s and above. In addition, the dynamics and loading of the muscle differed between the two modalities for most of the other variables we examined. The largest fiber length excursions were observed for the NHE, whereas the peak fiber lengths were greater for running at speeds of 6 m/s and above. The peak negative muscle-tendon unit power during the NHE was lower than during most of the running speeds we examined because the NHE is a slow exercise performed over approximately 4 s. Despite low negative muscle-tendon unit power during the NHE, the negative muscle-tendon unit work was significantly greater during the NHE compared to the flight phase of running at all speeds because of the long duration of the exercise. Our findings suggest that the two training modalities offer differing stimuli to the BFLH, and our results provide a baseline for future research on how the NHE and high-speed running mitigate injury risk.

We found that the NHE had a lower peak BFLH force compared to running at and above 7.5 m/s. These findings show that the NHE does not expose the BFLH to the force experienced during high-speed running, and, therefore, including high-speed running as part of hamstring injury prevention training guidelines is warranted as regular exposure may offer protection.^26^ Although running at and above 7.5 m/s the peak force was greater compared to during the NHE, no differences were found at 5, 6, and 7 m/s. In team-sports, such as soccer, it is common for athletes to perform a greater volume of running bouts at speeds between 5.5 and 7 m/s in training and matches,^42^ in which case our results show that the NHE provides a similar peak force exposure as to the most frequently encountered running speeds in team-sports. Even though the peak BFLH force during the NHE is lower compared to high-speed running, the stimulus provided appears to be sufficient to promote BFLH hypertrophy^43,44^ and increase hamstrings strength^16–19,21^, with these adaptations helping to lower the demands placed on the hamstrings during high-speed running.

Separate studies have used musculoskeletal modeling and simulation to characterize the BFLH force either during the NHE^30^ or high-speed running.^11–13^ Furthermore, a recent review^45^ concluded that the BFLH peak force is similar between the NHE and the flight phase of running at approximately 9 m/s based on the findings from separate studies.^11,30^ However, collectively interpreting the findings from separate studies is challenging due to differences in the musculoskeletal simulation methods and the cohort of recruited participants. In contrast, the findings from our study are not affected by those differences, as we used the same methods and participants for our investigation.

NHE and high-speed running training have been found to increase the length of BFLH fascicles,^16–18,20,21^, and this adaptation is regarded as a favorable adaptation to reduce injury risk.^16–20^ Fascicle lengthening is believed to arise from eccentric training modalities.^46^ We found that the muscle fibers operate at shorter lengths during the NHE compared to the flight phase of high-speed running (6 m/s and above), and fiber lengthening velocity was lower for the NHE compared to all running speeds. Taken together, these findings indicate that the NHE does not expose the BFLH to the fiber length and velocity demands of high-speed running. However, we also found that the fiber excursion was greater during the NHE in comparison to the flight phase of all running speeds. The results from NHE and high-speed running training interventions are equivocal in terms of which modality led to greater fascicle lengthening,^17,20^ and it is therefore not possible to deduce whether length or excursion is more important.

Muscle soreness is typically reported after NHE training,^47,48^ and the larger negative BFLH muscle-tendon unit work we found during the NHE may explain this. The larger negative BFLH muscle-tendon unit work during the NHE can be attributed to the longer duration of a repetition, which lasted on average 4 s, together with the greater fiber excursion, which contributes to muscle-tendon unit length excursion. Negative work as a stimulus for muscle adaptations has received limited attention; however, it warrants further investigation as it combines force and lengthening, which are important stimuli for muscle adaptations.^49^ Indeed, work on animals has shown that peak eccentric moment and strain are the two best predictors of sarcomerogenesis,^50^ and perhaps negative work, their composite, may aid to further explain muscle adaptations.

While our study has provided insights into the BFLH dynamics between the NHE and high-speed running, it is important to acknowledge several limitations. First, our participants had a variety of training backgrounds, and this heterogeneity may explain the variability in the outcome variables. Second, we found that the BFLH fiber lengthened slowly throughout the NHE, whereas ultrasound studies have shown that the BFLH fascicles only lengthen during the last 20% of the NHE.^30,51^ This difference may be due to Hill-type muscle models providing an estimate of uniform fiber length changes,^52^ whereas ultrasound enables localized fascicle length changes to be tracked.^53^ Third, we calibrated the muscle model parameters for each participant during their top running speed trial. Calibrating the muscle model parameters is essential for movements that require large joint moments to be generated, but we do not have independent measures of the muscle model parameters to test the accuracy of our calibrations. Future work should focus on developing a robust method for adapting muscle model parameters to an individual, enabling accurate and realistic simulations of athletic tasks. Finally, we have examined two common training modalities, and future studies could examine others.

## 5. Conclusion

When designing training programs, understanding the biomechanical demands from different training modalities is needed as the demands influence the adaptations of skeletal muscle in response to training. Our results revealed that the NHE and high-speed running provide dramatically different biomechanical demands. The NHE requires the hamstrings to generate a moderate level of force over a long duration as the muscle fiber lengthens very slowly. In contrast, high-speed running requires the hamstrings to develop a higher level of force at longer fiber lengths and higher fiber lengthening velocities over a very brief duration. To prevent hamstring strain injuries, it is unlikely that a single training modality will suffice. The differences we observed between the NHE and high-speed running suggest that these two training modalities offer complementary stimuli to promote favorable BFLH injury prevention adaptations.

## Acknowledgements

This work was supported by the National Science Foundation, the Stanford Graduate Fellowship in Science & Engineering, and the Joe and Clara Tsai Foundation through the Wu Tsai Human Performance Alliance. We thank Julie Muccini, Julie Kolesar, Tian Tan, and Sam Hamner for their support during the data collection sessions. We also thank Max Andrews, Glen Lichtwark, Dawit Lee, Jeff Stribling, and Alex Gonzalez for assistance.

## Data Statement

The data that support the findings of this study will be made openly available at https://simtk.org/projects/nhe_run_sims. The simulation framework used in this study will be made openly available at https://github.com/nicos1993/MTU_parameters_optimization.

