## Supplementary Materials for "Hamstrings muscle dynamics during the Nordic hamstring exercise and high-speed running"

*Marker locations:*

The markers were attached bilaterally on the anterior superior iliac spine, posterior superior iliac spine, iliac crest, lateral and medial femoral epicondyles, lateral and medial malleoli, calcaneus, second and fifth metatarsal heads, acromion, lateral and medial humeral epicondyles, and lateral and medial styloid processes, and unilaterally on the sternum and C7. An additional set of 20 tracking markers were attached to the thighs, shanks, upper arms, and lower arms. The medial femoral epicondyle and malleoli markers were removed prior to the NHE repetitions.

*Musculoskeletal model dynamics and modifications:*

For our simulations we used compliant Hill-type muscles. We formulated activation dynamics[1] explicitly with excitation as a control and activation as a state, and contraction dynamics[2] implicitly with normalized tendon force as a state and the derivative of normalized tendon force as a control. We also made several modifications to the Hill-type muscle model we used:[2] we widened the width of the active force-length curve by 15% to ensure we captured the operating region of a whole muscle as opposed to a single sarcomere,[3] we increased the maximum contraction velocity from 10 to 12 optimal fiber lengths per second based upon an in vivo estimate,[4] and we reduced the slope of the passive force-length curve by shifting the strain at maximum passive force from 0.7 to 0.8 to prevent an excessive passive force contribution at a normalized fiber length beyond optimal.

*Objective functions:*

We used the following objective functions for the calibration (1) and non-calibration (2) simulations:

$$w_a \sum_{n=1}^{N} \int_{t_i}^{t_f} a_n{}^2(t)\,dt + w_e \sum_{k=1}^{K} \int_{t_i}^{t_f} \frac{\left(e_k(t) - EMG_k(t)\right)^2}{EMG_{mapping}}\,dt + w_p \sum_{n=1}^{N} \int_{t_i}^{t_f} \tilde{f}_{PL_n}{}^2(t)\,dt$$
$$+ w_r \sum_{g=1}^{G} \int_{t_i}^{t_f} r_g{}^2(t)\,dt + w_d \sum_{n=1}^{N} \int_{t_i}^{t_f} \dot{\tilde{F}}_{T_n}{}^2(t)\,dt$$
$$+ w_m \sum_{i=1}^{I} \frac{\left(L_{o_i}^M - \hat{L}_{o_i}^M\right)^2}{\hat{L}_{o_i}^M} + w_m \sum_{i=1}^{I} \frac{\left(L_{S_i}^T - \hat{L}_{S_i}^T\right)^2}{\hat{L}_{S_i}^T} + w_m \sum_{i=1}^{I} \frac{\left(F_{o_i}^M - \hat{F}_{o_i}^M\right)^2}{\hat{F}_{o_i}^M} \quad (1)$$

$$w_a \sum_{n=1}^{N} \int_{t_i}^{t_f} a_n{}^2(t)\,dt + w_e \sum_{k=1}^{K} \int_{t_i}^{t_f} \frac{\left(e_k(t) - EMG_k(t)\right)^2}{EMG_{mapping}}\,dt + w_p \sum_{n=1}^{N} \int_{t_i}^{t_f} \tilde{f}_{PL_n}{}^2(t)\,dt$$
$$+ w_r \sum_{g=1}^{G} \int_{t_i}^{t_f} r_g{}^2(t)\,dt + w_d \sum_{n=1}^{N} \int_{t_i}^{t_f} \dot{\tilde{F}}_{T_n}{}^2(t)\,dt \quad (2)$$

where $a_n$ is the muscle activation of muscle $n$, $e_k$ is the muscle excitation of muscle $k$, $EMG_k$ is the EMG signal of muscle $k$, $EMG_{mapping}$ corresponds to the number of muscle excitations sharing a given EMG signal, $\tilde{f}_{PL_n}$ is the passive force-length multiplier of muscle $n$, $r_g$ is the reserve joint moment actuator of joint $g$, $\dot{\tilde{F}}_{T_n}$ is the derivative of normalized tendon force of muscle $n$, $L_{o_i}^M$ is the optimal fiber length of muscle $i$, $L_{S_i}^T$ is the tendon slack length of muscle $i$, $F_{o_i}^M$ is the maximal isometric force of muscle $i$, $\hat{L}_{o_i}^M$ is the nominal optimal fiber length of muscle $i$, $\hat{L}_{S_i}^T$ is the nominal tendon slack length of muscle $i$, and $\hat{F}_{o_i}^M$ is the nominal maximal isometric force of muscle $i$. $N$ and $G$ are the total number of muscles and reserve actuators, respectively, $K$ is the total number of muscle excitation-EMG tracking terms, and $I$ are the total number of muscles for a single limb. $t_i$ and $t_f$ are the initial and final times, respectively, of each flight phase or NHE repetition. $w_a$, $w_e$, $w_p$, $w_r$, $w_d$, and $w_m$ are the weights of the terms in the objective functions. We set the weights as follows: $w_a$= 0.1, $w_e$ = 5, $w_p$ = $10^{-4}$, $w_r$ = $10^3$, $w_d$ = 1, $w_m$ = 0.01.

*Constraints:*

For all simulations, we included differential constraints to enforce muscle activation and contraction dynamics, and algebraic path constraints to enforce muscle-tendon force equilibrium and ensure that the net muscle moment plus reserve actuator matched the net joint moment. The joint moment constraints were imposed for the net joint moments corresponding to hip flexion-extension and abduction-adduction, knee flexion-extension, and ankle plantarflexion-dorsiflexion for both limbs. We also included inequality constraints to limit normalized fiber lengths to be between 0.4 and 1.8.

For the calibration simulation we also included two additional muscle-tendon force equilibrium constraints per muscle. These constraints were imposed at the minimum and maximum muscle-tendon unit lengths for a given muscle, extracted from across a given participant's entire set of experimental trials. For these constraints, we set muscle-tendon unit velocity to zero and muscle activation to 1. We simultaneously constrained normalized fiber lengths to fall between 0.4 and 1.8. The inclusion of these additional constraints for the calibration simulation prevented infeasible muscle kinematics states from being reached in the subsequent simulations. In addition, to prevent unrealistic combinations of maximal isometric forces between muscles when calibrating, we imposed constraints to ensure they all scaled uniformly.

*Bounds:*

We set the lower and upper bounds for muscle excitations $\boldsymbol{e}(t)$, muscle activations $\boldsymbol{a}(t)$, normalized tendon forces $\tilde{\boldsymbol{F}}_{\boldsymbol{T}}(t)$, the derivative of normalized tendon forces $\dot{\tilde{\boldsymbol{F}}}_{\boldsymbol{T}}(t)$, and reserve actuators $\boldsymbol{r}(t)$ as follows:

$$\mathbf{0} \leq \boldsymbol{e}(t) \leq \mathbf{1}, \tag{3}$$

$$\mathbf{0} \leq \boldsymbol{a}(t) \leq \mathbf{1}, \tag{4}$$

$$\mathbf{0} \leq \tilde{\boldsymbol{F}}_{\boldsymbol{T}}(t) \leq \mathbf{5}, \tag{5}$$

$$-\mathbf{100} \leq \dot{\tilde{\boldsymbol{F}}}_{\boldsymbol{T}}(t) \leq \mathbf{100}, \tag{6}$$

$$-\mathbf{50} \leq \boldsymbol{r}(t) \leq \mathbf{50} \tag{7}$$

where the bounds for $\tilde{\boldsymbol{F}}_{\boldsymbol{T}}(t)$ and $\dot{\tilde{\boldsymbol{F}}}_{\boldsymbol{T}}(t)$ were set based on the work of Falisse et al.[5] The bounds for each optimal fiber length $L_{o_i}^M$ were set using the optimal fiber length mean and standard deviation reported by Son et al.[6] if possible:

$$\mu_{L_{o_i}^M} - 3\sigma_{L_{o_i}^M} \leq L_{o_i}^M \leq \mu_{L_{o_i}^M} + 3\sigma_{L_{o_i}^M} \tag{9}$$

Otherwise, the lower and upper bounds were set to ± 15% of the nominal optimal fiber length $\hat{L}_{o_i}^M$ from scaling the generic model to each participant. The lower and upper bounds for each tendon slack length $L_{S_i}^T$ were set to ± 25% of the nominal tendon slack length $\hat{L}_{S_i}^T$ post model scaling, and follows the bounds set by Bianco et al.[7] Lastly, the lower and upper bounds for each maximal isometric force $F_{o_i}^M$ were set to 50% and 250%, respectively, of the nominal maximal isometric force $\hat{F}_{o_i}^M$ in the generic model. The upper bounds for each $F_{o_i}^M$ were necessary to avoid reliance on reserve actuators and are in accordance with the increases used by Swinnen et al.[8] and Van Hooren et al.[9] to perform inverse simulations of running at 3.33 m/s and hamstrings training exercises, respectively.

*Initial guess:*

We set the initial guess for each type of state and control variable as a constant value across the discretized mesh: 0.1 for $\boldsymbol{e}(t)$ and $\boldsymbol{a}(t)$, 0.3 for $\tilde{\boldsymbol{F}}_{\boldsymbol{T}}(t)$, 5 for $\dot{\tilde{\boldsymbol{F}}}_{\boldsymbol{T}}(t)$, and 0 for $\boldsymbol{r}(t)$. We applied a different initial guess to achieve convergence for one participant's set of simulations. This featured drawing a different single sample for $\boldsymbol{e}(t)$, $\boldsymbol{a}(t)$, $\tilde{\boldsymbol{F}}_{\boldsymbol{T}}(t)$ and $\dot{\tilde{\boldsymbol{F}}}_{\boldsymbol{T}}(t)$ from a uniform distribution and then applying the randomly drawn sample as a constant value across the discretized mesh. The uniform distribution was bounded between 0.1 and 0.8 for $\boldsymbol{e}(t)$, $\boldsymbol{a}(t)$ and $\tilde{\boldsymbol{F}}_{\boldsymbol{T}}(t)$, and between -0.4 and 0.4 for $\dot{\tilde{\boldsymbol{F}}}_{\boldsymbol{T}}(t)$. We continued to initialize $\boldsymbol{r}(t)$ as zero for this initial guess. For the calibration simulations, we initialized $\boldsymbol{L}_{\boldsymbol{o}}^{\boldsymbol{M}}$, $\boldsymbol{L}_{\boldsymbol{S}}^{\boldsymbol{T}}$, and $\boldsymbol{F}_{\boldsymbol{o}}^{\boldsymbol{M}}$ to their nominal values.

*Additional muscle excitations and EMG signals:*

The patterns of the muscle excitations for the vastus medialis and lateralis closely matched the corresponding measured EMG signals for both running and the NHE (Figure S1), with a maximum root mean square error (RMSE) of 0.03. EMG tracking of the soleus, tibialis anterior, and medial gastrocnemius resulted in higher RMSEs than the other muscles. The largest RMSE was for the medial gastrocnemius during the flight phase of running at 8 m/s.

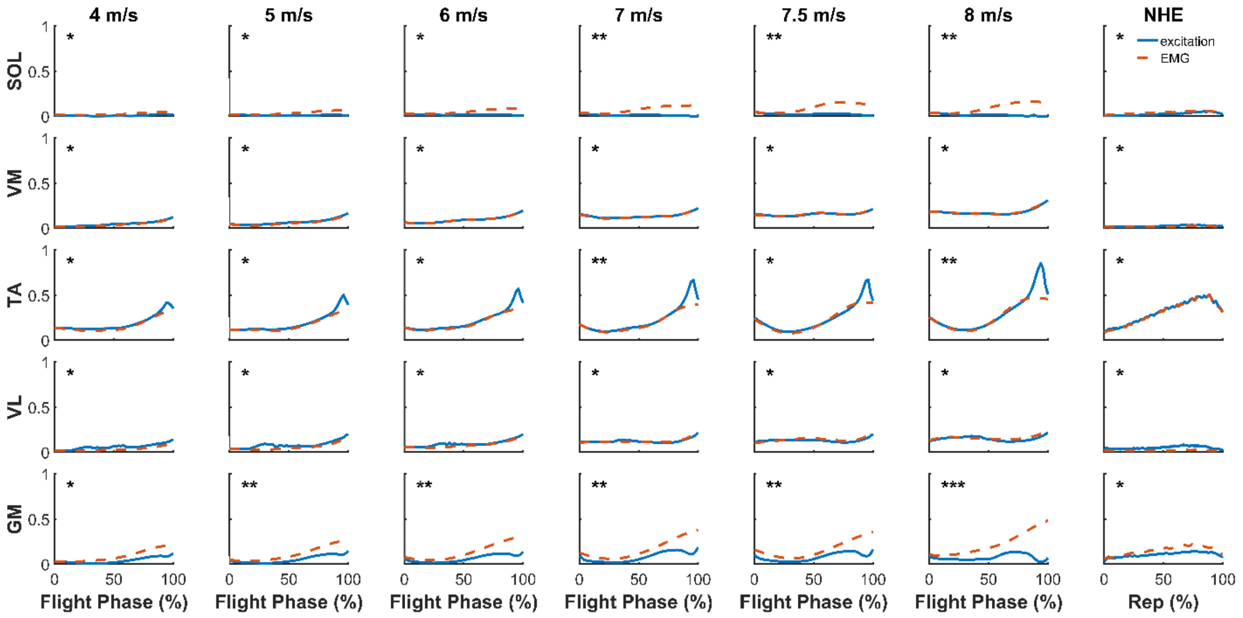


**Figure S1** – Simulated muscle excitations (solid blue line) and processed EMG signals (dashed red line) during the flight phase of high-speed running at different speeds and during the NHE. Each curve is the mean from across all participants' trials. Asterisks indicate the magnitude of the root mean square error (RMSE) between the mean muscle excitations and processed EMG signals: * = RMSE of ≤ 0.06, ** = 0.06 < RMSE ≤ 0.12, and *** = 0.12 < RMSE ≤ 0.20. The figure includes a subset of key lower-limb muscles, including the soleus (SOL), vastus medialis (VM), tibialis anterior (TA), vastus lateralis (VL), and medial gastrocnemius (GM).

**Supplementary Materials References**
